# GenAPI: a tool for gene absence-presence identification in fragmented bacterial genome sequences

**DOI:** 10.1101/658476

**Authors:** Migle Gabrielaite, Rasmus L. Marvig

## Abstract

Bacterial gene loss and acquisition is a well-known phenomenon which contributes to bacterial adaptation through changes in important phenotypes such as virulence, antibiotic resistance and metabolic capability. While advances in DNA sequencing have accelerated our ability to generate short-read genome sequencing to disentangle phenotypic changes caused by gene loss and acquisition, the short-read genome sequencing often results in fragmented genome assemblies as a basis for identification of gene loss and acquisition events. However, sensitive and precise determination of gene content change for fragmented genome assemblies remain challenging as analysis needs to account for cases when only a fragment of the gene is assembled or when the gene assembly is split in more than one contig.

We developed GenAPI, a command-line tool that is designed to compare the gene content of bacterial genomes for which only fragmented genome assemblies are available. GenAPI, unlike other available tools of similar purpose, accounts for genome assembly imperfections and aims to compensate for them. We tested the performance of GenAPI on three different datasets to show that GenAPI has high sensitivity while it maintains precision when dealing with partly assembled genes in both simulated and real datasets. Furthermore, we compared and evaluated the performance of GenAPI with six popular tools for gene presence-absence identification. While we find that the compared tools have the same precision and recall rates when analyzing complete genome sequences, GenAPI performs better than the other tools on fragmented genome assemblies.

## Background

Finding differences in the gene repertoire between bacterial clones is important to understand the genetic basis of differences in phenotypes such as virulence, antibiotic resistance, and metabolic capability. Also, phylogenetic analysis including information about the absence or presence of genes helps to inform about the pace and mechanisms by which genes are lost and acquired in bacterial populations [1]. Thus, genome-wide analysis of the gene presence or absence is necessary for a better understanding of bacterial evolution and adaptation.

There exist multiple open-source bioinformatics tools available for gene presence-absence identification. Each tool has its own set of advantages and limitations. Here, we focus on tools that use assembled genomes as input. Some tools, e.g. PanSeq [2], are based on alignment of the query genome sequence to a reference genome to specifically test for the presence-absence of genes that are present in the given reference. Accordingly, this approach is limited knowing that prophage and plasmid genes constitute a major part of the variable bacterial genetic content [3]. Other tools, e.g. Roary [4], SaturnV [5], PanDelos [6], panX [7], BPGA [8] and EDGAR [9] construct a pan-genome from inputted genome assemblies and then determine the gene set that is present in each of the genome assemblies. The performance of both approaches depends on the completeness of the queried genome sequences, and all previously mentioned tools are designed for analysis of near-complete genomes with only minor parts of genes missing due to sequencing and assembly imperfections. As a result, there is a need for analytical tools that are designed to account for highly fragmented genome assemblies (often the product of *de novo* genome assemblies based on short-read sequence data where genome assemblies have tens to hundreds of contigs) that else would result in a large number of false calls for gene being absent in the assembly.

Here, we introduce GenAPI, a command line tool for identification and comparison of the gene content in bacterial clones of the same species. We specifically developed GenAPI to work successfully on fragmented genome assemblies to account for cases when only a fragment of the gene is assembled or when the gene assembly is split in more than one contig (Figure 1). The compensation for imperfections of sequencing and assembly proves to minimize the false gene absence calls while maintaining the true gene absence call rates.

**Figure 1.**
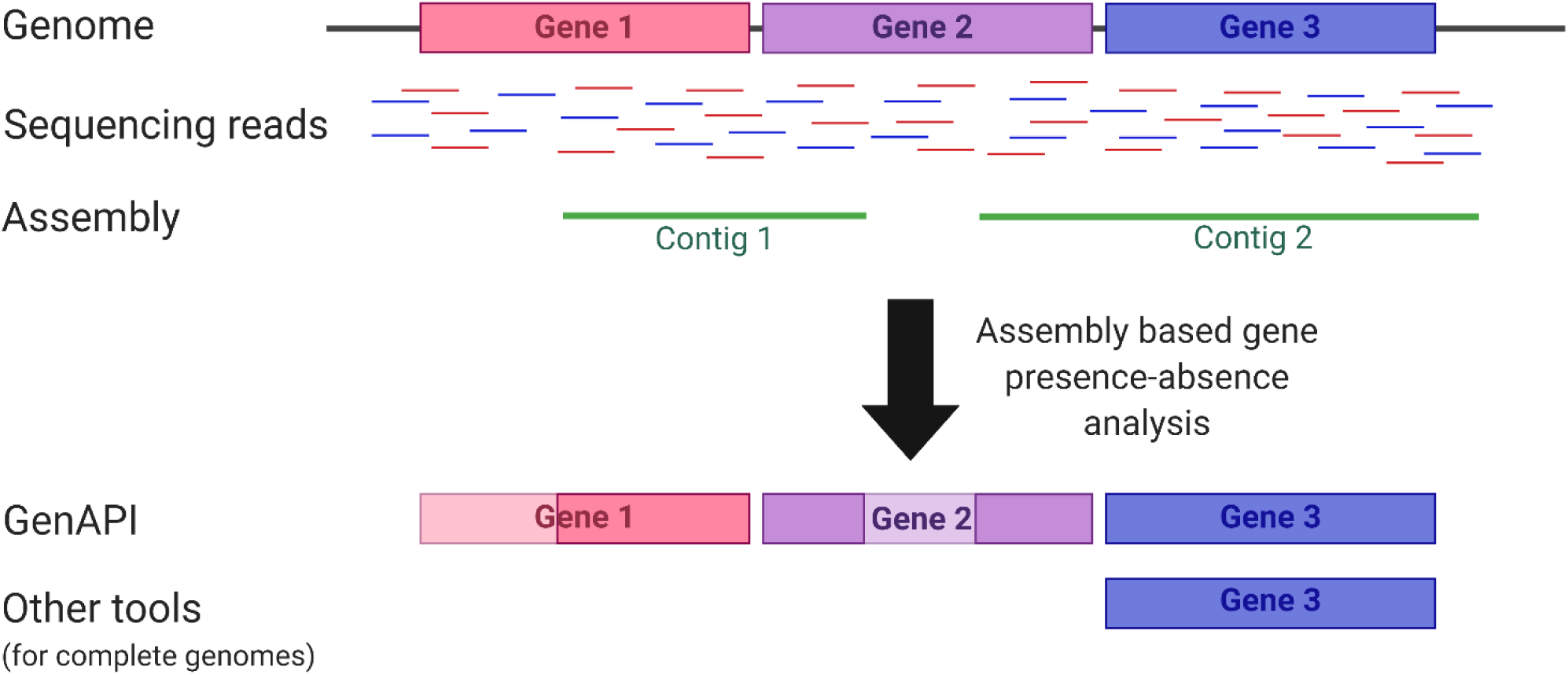
Schematic visualization of *de novo* assembly imperfections which are addressed by GenAPI to improve the gene presence-absence identification.

## Results

### Description of GenAPI algorithm

GenAPI is designed to run in a Unix environment and requires BLAST+, CD-HIT and Bedtools softwares [10, 11, 12]. R with pheatmap library [13, 14] and RAxML [15] softwares are needed for optional gene presence-absence matrix visualization and phylogenetic tree generation. The step-wise workflow of GenAPI is shown in Figure 2A where annotated genome sequence files are taken as an input and the pan-genome (the total gene repertoire of the bacterial clones [16]) is generated by clustering the genes with CD-HIT-EST with 90% identity and 80% sequence length overlap requirements. The 80% sequence overlap requirement allows incompletely sequenced or assembled genes to be clustered correctly. Gene clustering similarity thresholds should be reconsidered if GenAPI is used on non-closely related bacterial genomes. The best alignments between genes of the pan-genome and each genome sequence are identified by performing all-vs-all sequence comparison. Nucleotide alignments were chosen to be used over amino acid alignments as it enables more specific evaluation of sequence identity (i.e., one amino acid can be defined by multiple codons). Finally, the genes are defined as present or absent according to the user-provided thresholds (default: gene is present if best alignments have minimum 25% coverage with 98% identity or 50% coverage with 90% identity). More detailed method explanation is provided in Supp. Figure 1.

**Figure 2.**
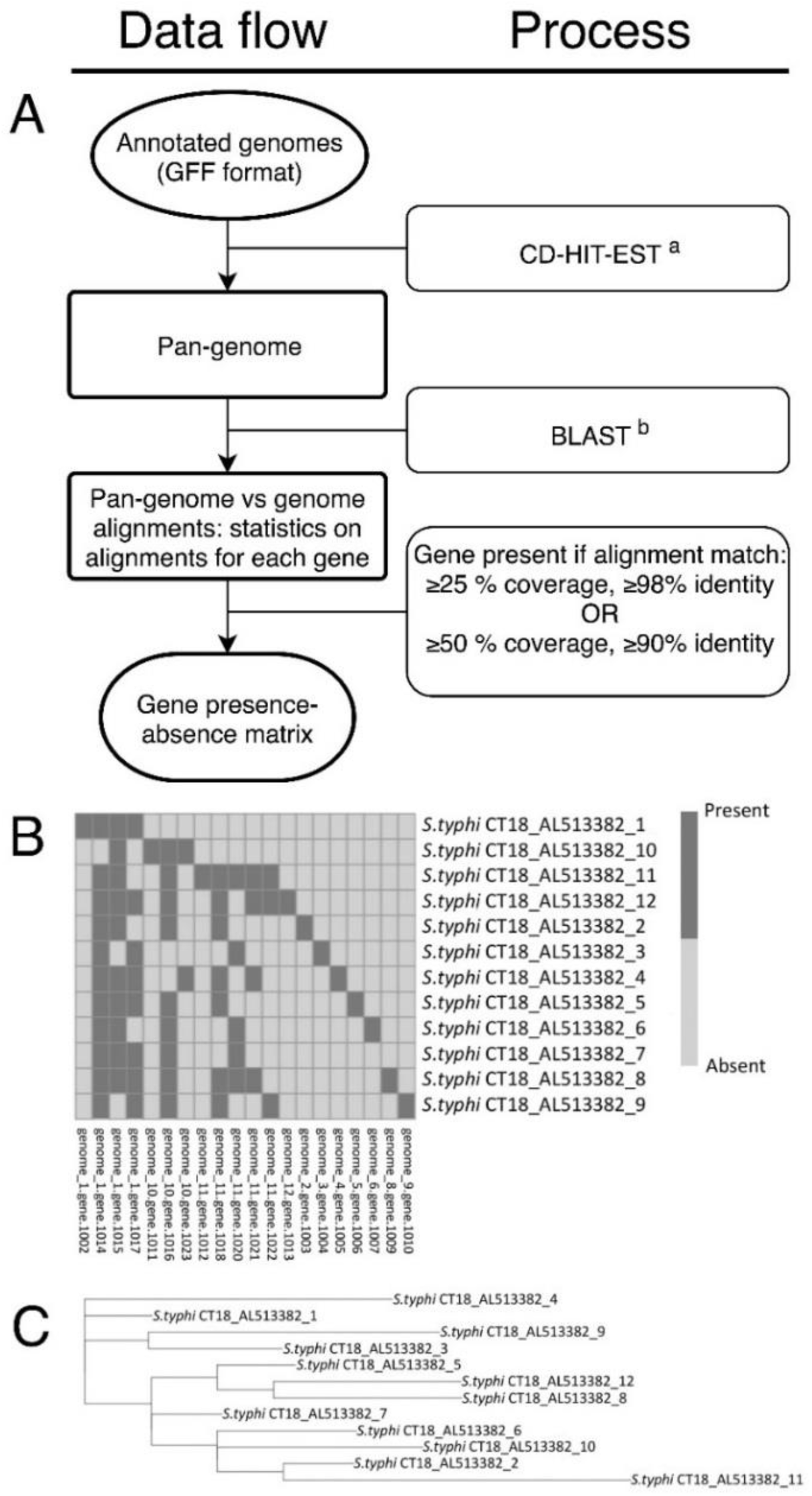
GenAPI workflow. (A) GenAPI data flow including the tools [17]^a^, [18]^b^ or algorithms used in each step. GenAPI outputs (B) gene presence-absence matrix and (C) and phylogenetic tree based on gene presence-absence.

The gene is defined as present even if it is only partly aligned but with high identity, this allows to compensate for sequencing and assembly imperfections while not compromising precision. The requirement for only 25% alignment coverage with nearly perfect identity is based on low likelihood of having a random, high-identity average length alignment (Figure 1). Nonetheless, genes shorter than 150 base pairs (bp) are recommended to be excluded from the gene presence-absence analysis as alignment of short gene sequences may produce unspecific alignments that may pass threshold for defining a gene as present (e.g. a 100 bp gene would be defined as present with an unspecific 25 bp alignment). Short genes are excluded from the analysis by default; however, this setting can be changed or turned off if needed. Furthermore, GenAPI is developed for identification of gene sequences that are completely lost or acquired in the genome, therefore it will not identify copy number changes of multicopy genes (including pseudogenes) or partial gene deletions. To address these issues a long-read whole genome sequencing is recommended. [17]

GenAPI outputs several files: (1) a pan-genome file, (2) a gene presence-absence matrix with information on the presence of each gene in each of analyzed genome sequences, and (3) a list of best alignment statistics for each gene in each analyzed genome sequence. Furthermore, a heatmap visualization of gene presence-absence across all genomes and a maximum-likelihood phylogenetic tree based on gene presence-absence information is implemented in GenAPI (Figure 2B-C).

### Evaluation of GenAPI and comparison with other tools for gene presence-absence identification

We evaluated the performance of GenAPI on three test datasets with genome assemblies: two simulated datasets with *in silico* introduced variation in gene content, and one real dataset with known gene deletions. Furthermore, we compared the performance of GenAPI against Roary, SaturnV, PanDelos, BPGA, panX and EDGAR which are tools developed for similar but not identical purposes. None of the tools which we compared against were developed for fragmented genome assemblies; however, fragmented genome assemblies are most often the output from genome sequencing experiments and there are no available tools developed for gene presence-absence analysis in fragmented genome assemblies.

If no insertions/deletions are introduced to the genomes, all genes in all analyzed genomes will be present, therefore, only gene absence was measured to evaluate the tools’ performance. The results of the predicted gene absence of the recall, precision and F1 scores are shown in Table 1. F1 measure was calculated in order to summarize precision and recall scores as it is a harmonic average of these two measures. Recall, precision and F1 scores were calculated by using the following formulas:

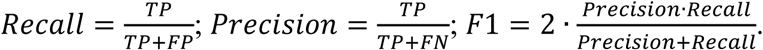

**Table 1.** Recall, precision and F1 score of GenAPI, panX, BPGA, Roary, PanDelos, EDGAR and SaturnV using simulated and real datasets.

|  |  | GenAPI | panX | BPGA | Roary | PanDelos | EDGAR | SaturnV |
| --- | --- | --- | --- | --- | --- | --- | --- | --- |
| *S. typhi*<sup>a</sup> | Precision | 1 | 1 | 0.93 | 1 | NA | 1 | 1 |
|  | Recall | 1 | 1 | 1 | 1 | NA | 1 | 1 |
|  | F1 | 1 | 1 | 0.97 | 1 | NA | 1 | 1 |
| *P. aeruginosa*<sup>b</sup> | Precision | 0.94 | 0.37 | 0.31 | 0.31 | 0.45 | 0.15 | 0.11 |
|  | Recall | 1 | 1 | 0.94 | 1 | 1 | 1 | 1 |
|  | F1 | 0.97 | 0.54 | 0.46 | 0.47 | 0.62 | 0.26 | 0.20 |
| *E. coli*<sup>c</sup> | Precision | 0.95 | 0.47 | 0.26 | 0.23 | 0.24 | 0.12 | 0.09 |
|  | Recall | 0.98 | 1 | 0.88 | 1 | 0.71 | 1 | 1 |
|  | F1 | 0.97 | 0.64 | 0.40 | 0.38 | 0.36 | 0.21 | 0.17 |

a - simulated *S. typhi* dataset with complete genome sequences used in Roary publication [4]; b - simulated *P. aeruginosa* dataset with partly assembled gene instances; c – real *E. coli* dataset [18].

For simulated *P. aeruginosa* and long-term *E. coli* experiment [18] datasets Prokka version 1.11 [19] was used for gene annotation. Sequencing reads from both datasets were assembled using SPAdes version 3.10.1 software with default parameters [20]. *P. aeruginosa* and *E. coli* assembly statistics are reported in Supp. Table 1 and Supp. Table 2, respectively. Information about known deletions in *E. coli* experiment dataset was obtained from Barrick *et al*. (2009) [18] study. *S. typhi* dataset was preprocessed as described by Page *et al*. (2015) [4]. All tools were tested with default parameters, with an exception for Roary for which paralog splitting was disabled since the other tools do not split identical sequences (paralogs). Default parameters were chosen assuming that they are the most optimal parameters for the tool. Sequencing reads for *P. aeruginosa* simulated dataset were simulated using ART software [21] with 150bp paired end sequences with 200bp insert sizes and 20x average genome coverage.

First, all tools except PanDelos were tested on the same simulated *Salmonella typhi* dataset that was used for the evaluation of Roary in its own publication [4]. PanDelos was excluded from the analysis as the tool did not finish the analysis in 24 hours since its start. The six included tools identified all 181 instances where genes were absent from genomes (GenAPI did not include one gene as it was shorter than the default 150bp gene length requirement) (Supp. Table 3). BPGA additionally made 12 false calls of gene being absent. The other tools did not falsely call any genes to be absent.

Second, we simulated sequencing reads for a dataset of 8 *P. aeruginosa* genome sequences with known deletions in order to test the tools on data representing fragmented genome assemblies (available at https://github.com/MigleSur/GenAPI/tree/master/test_samples). In total, there were 49 deleted genes (Supp. Table 5). All tools except BPGA, which missed 3 deletions, correctly identified all 49 absent genes. However, false positive deletion calls were made by all tools. 145, 387 and 103 genes were false positively called as absent by Roary, SaturnV and BPGA, respectively. PanDelos and panX performance was better and resulted in 60 and 83 genes to be false positively called as deleted, respectively. On the other hand, only 3 false positive deletions were predicted by GenAPI (Supp. Table 3).

Finally, we tested the tools on a dataset of 6 genome sequences with 102 known deletions from the *E. coli* long-term evolution experiment performed by Richard Lenski [18]. Roary, SaturnV, EDGAR and PanX called all true deletions, and GenAPI closely followed these tools except missing two copy number changes of homologous prophage genes. BPGA and PanDelos failed to identify 12 and 30 true deletions, respectively. The number of false positive calls of gene deletions was observed to be high for Roary, SaturnV, panX, PanDelos, BPGA and EDGAR (115-1009 false positive deletion calls) in contrast to GenAPI (5 false positive calls) (Supp. Table 3). F1 scores from Table 1 show a summary of how well each tool performs with each of the datasets.

## Discussion

While there exist tools for bacterial gene content identification and comparison, our developed tool—GenAPI—was specifically designed for the analysis of fragmented, closely related genome sequences. Fragmented genome sequences are often the product of *de novo* genome assembly based on short-read sequence data, and GenAPI will therefore be suitable for use in multiple studies including bacterial genome sequence data.

Here we tested the performance of GenAPI on three different datasets to show that GenAPI has a high sensitivity while it maintains precision when dealing with partly assembled genes in both simulated and real datasets. While all tested tools except BPGA and PanDelos are excellent at identifying true gene absence (i.e. high sensitivity), only GenAPI has a low rate of false positive gene absence calls in fragmented genomes (i.e. high precision). The higher precision of GenAPI could be explained by the lower requirement for sequence alignment length which is conditional to a high alignment identity. The other tools included in the study have high requirements for sequence alignment length which does not cause any false gene absence predictions in simulated complete genome dataset (*S. typhi* dataset); however, it markedly affects the precision when applied to fragmented simulated (*P. aeruginosa*) and real (*E. coli*) datasets (Table 1, Supp. Table 3).

We have shown that none of the other popular gene presence-absence identification tools are well suited for analysis of fragmented genome assemblies and the performance of GenAPI is better when analyzing fragmented genomes.

## Author statements

### Data Summary

*S. typhi* simulated sequence data was previously published by Page *et al*. (2015);*P. aeruginosa* simulated reads are available from: https://github.com/MigleSur/GenAPI/tree/; *E. coli* sequencing data was published by Barrick *et al*. (2009) and are publicly available in Sequence Read Archive under accession SRP001369.

### Authors and contributions

MG and RLM designed the methodology, MG developed the tool, MG and RLM analyzed and interpreted the results. MG prepared the original draft, MG and RLM were involved in revising, editing the draft, reading and approving the final manuscript.

## Acknowledgements

We thank Anders Krogh for consultations and help during GenAPI development. Figure 1 was created using BioRender (https://biorender.com/).

